# Causal inference on microbiome-metabolome relations via *in silico in vivo* association pattern analyses

**DOI:** 10.1101/2021.03.15.435397

**Authors:** Johannes Hertel, Almut Heinken, Ines Thiele

## Abstract

The effects of the microbiome on the host’s metabolism are core to understanding the role of the microbiome in health and disease. Herein, we develop the paradigm of *in silico in vivo* association pattern analyses, entailing a methodology to combine microbiome metabolome association studies with *in silico* constraint-based microbial community modelling. By dissecting confounding and causal paths, we show that *in silico in vivo* association pattern analyses allows for causal inference on microbiome-metabolome relations in observational data. Then, we demonstrate the feasibility and validity of our approach on a published multi-omics dataset (n=346), demonstrating causal microbiome-metabolite relations for 43 out of 53 metabolites from faeces. Finally, we utilise the identified *in silico in vivo* association pattern to estimate the microbial component of the faecal metabolome, revealing that the retrieved metabolite prediction scores correlate with the measured metabolite concentrations, and they also reflect the multivariate structure of the faecal metabolome. Concluding, we integrate with hypothesis free screening association studies and knowledge-based *in silico* modelling two major paradigms of systems biology, generating a promising new paradigm for causal inference in metabolic host-microbe interactions.

## Introduction

The determination of the microbiome’s metabolic functions is a key challenge in understanding the contribution of the gut microbiome to health and disease [1–3]. As metabolic functions are shared across phylogenetic classes [4], differences in composition do not necessarily translate in differences in metabolic output. Therefore, analyses of the microbiome composition alone cannot give conclusive insights into the collective metabolic output of a community. In the light of this challenge, researchers have repeatedly tried to shine a light on metabolic functions of microbes via integrating microbial abundance data with metabolome data via statistical association studies [5–8]. Especially faecal metabolomics, being closest to a direct functional readout, has been used for statistical screening for associations [6, 8, 9]. However, statistical screenings can easily result in false positives [10], and recent modelling has shown that microbe-metabolite associations are prone to be the result of confounding, especially due to the multivariate nature of both types of omics datasets [11]. Moreover, from the viewpoint of causal statistics, extracting causal models correctly from observational data requires full information on all relevant confounding variables [12]. However, measuring and conceptualising all relevant confounders poses conceptual and practical problems for a concrete microbiome-metabolome study, partly due to limited knowledge on relevant confounding factors [13]. Hence, results of statistical hypothesis free screening approaches are often difficult to interpret and to embed into the knowledge already gathered about microbial biology [13, 14].

Integrating the genetic content of the microbial community with knowledge of microbial biology while respecting basic laws of nature, such as conservation of mass and charge, constraint-based modelling and reconstruction analysis (COBRA) [15] allows a fine-graded mapping of metabolic functions of microbial communities [16]. Thus, COBRA is optimally suited to complement microbiome-metabolome association studies by delivering biological context in a quantitative way [17]. COBRA community models offer quantifications of the feasible range of metabolic fluxes given a diet, allowing consequently the calculation of, for instance, the metabolite secretion potential into a simulated faecal compartment of a given microbial community *in silico* [16]. In contrast to species metabolome association analysis, COBRA microbial community modelling allows the direct deterministic calculation of the contribution of a species to the output of the whole community [17]. As such, COBRA modelling results are not impacted in the same way as statistical associations by confounding caused by physiological and behavioural attributes of the host. However, while COBRA microbial community modelling has already been applied to investigate metabolic functions in Parkinson’s disease [18, 19] and inflammatory bowel disease [17], the predictions of the COBRA microbial community models have not been integrated systematically with metabolomic *in vivo* data to validate the predicted metabolic functions.

Here, we develop first the theoretical frameworks to combine COBRA microbial community modelling with microbiome metabolome association studies, outlining the causal and confounding paths effective in species-metabolite associations *in vivo* and *in silico*. Building on these theoretical considerations rooted in causal inference theory, we develop a methodological paradigm, which we call ‘*in silico in vivo* association pattern analyses’, allowing for causal inference on microbiome-metabolite relations in theory. Using published metagenomic data in conjunction with faecal metabolome data from [8], we then demonstrate that community modelling systematically predicts the statistical associations pattern between microbiome measurements and faecal metabolome measurements, justifying the proposed methodology. Finally, we show that our framework can be used to derive predictions scores for faecal metabolite concentrations from microbiome composition data, characterising the microbial component of the faecal metabolome. Our work highlights how metabolomics and metagenomics in combination with COBRA microbial community modelling can be utilised to improve mechanistic understanding, generate hypotheses, and identify and validate biomarkers for metabolic functions in human health and disease.

## Results

### Theoretical frameworks

The challenge presented by *in vivo* species-metabolite association studies, especially in observational data, is to disentangle the various sources of correlation. To this end, we classify the various sources of correlation between species and faecal metabolite concentrations *in vivo*. In a second step, we examine how these sources of correlation influence *in silico* species-flux associations derived from COBRA community models. Finally, to integrate species-metabolite association studies utilising faecal metabolome data with COBRA modelling, we introduce herein a theoretical framework of how *in silico* calculations refer to *in vivo* correlations, resulting in an analysis paradigm that allows for causal inference on metabolome-microbiome relations.

#### Causal and confounding paths in species-metabolite associations in vivo

Statistical correlation between species abundances and metabolite concentrations in observational data can result either of confounding or causation. We discuss first the causal paths by utilising directed acyclic graphs [12] leading to species-metabolite association by physiological, biochemical, or ecological mechanisms (Fig 1). First, a species can produce or consume a metabolite, directly influencing the metabolite’s concentration (direct metabolic causation). Second, a species may produce an intermediate, which is then converted by another microbe into the metabolite under consideration (indirect metabolic causation); an effect, which can, for example, be seen in microbial bile acid metabolism [17]. Third, a microbe can influence the abundance of another microbe via competition or cooperation [20], which in return is causally linked to the metabolite (ecological causation). Fourth, a microbe may modulate a physiological factor of the host (for example, inflammation [21]), which in return may influence the concentration of the metabolite under consideration. Fifth, a microbe may also modulate a behavioural factor, for example, dietary habits [22], which then impacts the metabolite’s concentration (behavioural causation) (Fig 1).

**Figure 1:**
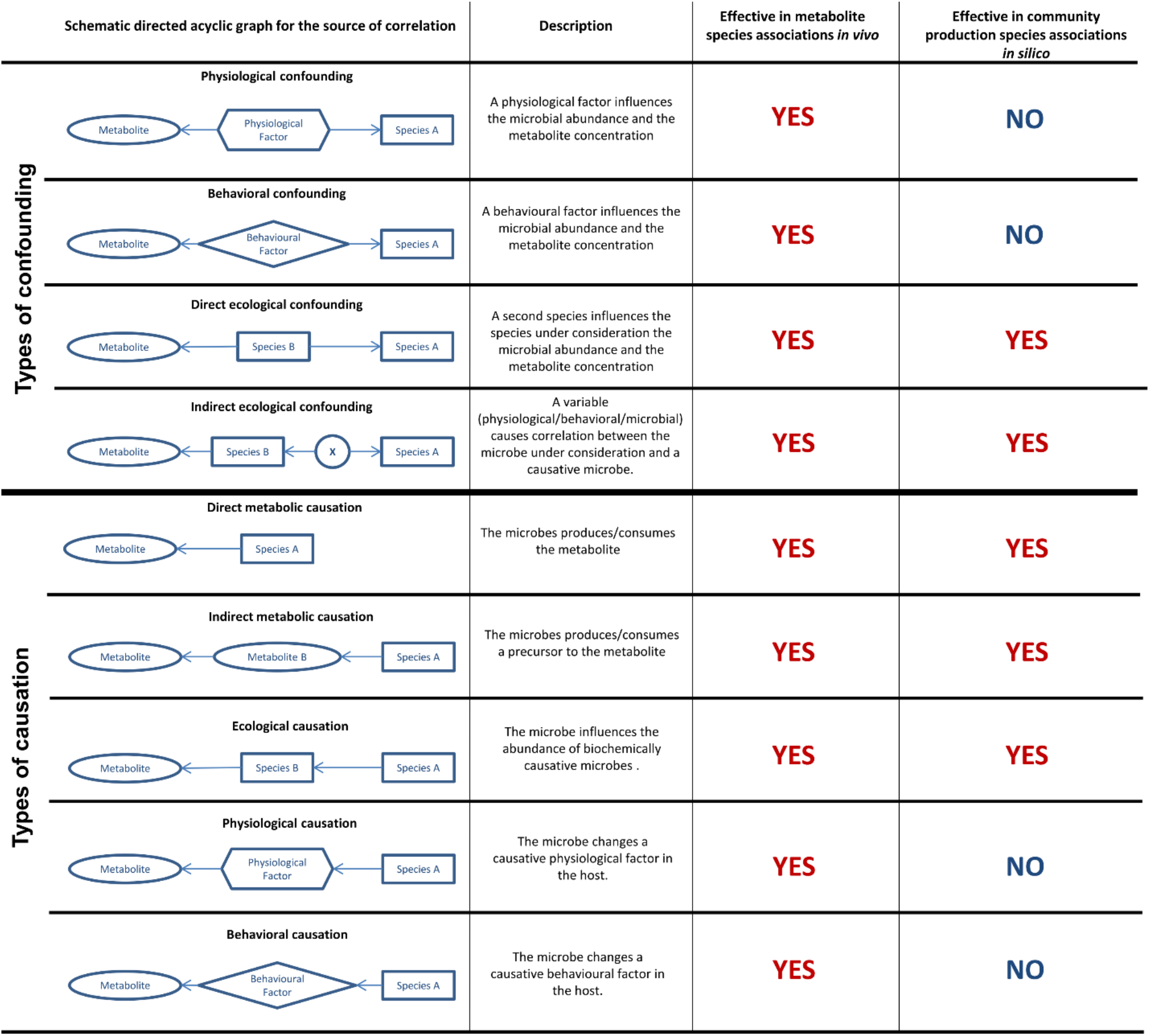
Causal and confounding paths in *in vivo* and *in silico* species metabolite association studies

However, confounding plays an equally important role as a source of correlation (Fig 1). Physiological factors of the host, for example, constipation [23], may impact the abundance of a microbial species, while also affecting a metabolite (physiological confounding), inducing correlation between the species abundance and the metabolite, which are unrelated otherwise. Second, behavioural traits of the host, for example, diet [24], may also influence microbial abundances and metabolite concentrations (behavioural confounding). Additionally, another species may be causally related to the species of interest, while impacting the levels of a metabolite (direct ecological confounding). Alternatively, a third factor (microbial, physiological, or behavioural) may induce correlation between the species under consideration and another species, which is causally linked to the metabolite’s concentration (indirect ecological confounding). In all these scenarios of confounding, we would observe a correlation between species abundance and metabolite concentrations without any underlying causal relation.

When determining associations by integrating metabolomic and metagenomic data, the various causal and confounding paths are added up to one single association statistic (e.g., the regression coefficient), making statistically significant associations difficult to interpret. Therefore, we need to integrate further information into the interpretation of species-metabolite association to allow for causal inference. Next, we show that COBRA community models can provide the necessary, additional context on the nature of metabolite-species associations to allow for a more refined interpretation of association statistics.

#### Causal and confounding paths in species-metabolite associations in silico

An individual COBRA microbial community model provides the metabolic secretion profile of the microbial community under consideration by calculating maximal net production fluxes from a set of diet constraints [16], the underlying genome scale reconstructions of the individual microbes, and the measured composition of the microbial community. If we have a population of microbial communities and their corresponding COBRA microbial community models, we can derive an *in silico* flux-species correlation pattern by correlating the species abundances with the overall community net metabolite production capacities. These association patterns, expressed in fluxes rather than concentrations, can be seen as theoretical counterparts to the *in vivo* species metabolite association patterns from metabolome-microbiome association studies. Importantly, as multiple studies have shown, variance in microbial abundance, and thereby variance in metabolic secretion and consumption, influence the metabolic profiles of the host [6, 8, 9]. Thus, if COBRA microbial community models are valid descriptions of the actual microbial activity, then the variation in the *in silico* net secretion pattern translates into *in vivo* variation in metabolite concentrations in the host. However, this assumption needs further theoretical considerations, as COBRA microbial community models do not reflect all causal and confounding paths effective in *in vivo* species-metabolite associations (Fig 1).

COBRA microbial community modelling allows calculating the direct contribution of a species to the metabolic net production profile of a community, thereby quantifying the direct and indirect metabolic causal effects [17]. Additionally, it allows for quantifying the ecological effects as well, although no inference on the nature (i.e., causal vs. confounding) of the ecological effects can be made from community modelling alone [25]. In essence, all confounding and causal pathways, which lead to correlation among species abundances, impact the output of COBRA microbial community models, explicitly including ecological causality and the two types of ecological confounding (Fig 1). Crucially, if the diet is held constant across the interrogated population of the computational microbial community models as done in [18, 19], *in silico* flux-species associations are independent of the concrete physiological or behavioural attributes of the host, which means that neither physiological, or respectively behavioural, confounding nor causality are represented. In conclusion, for *in silico* species-metabolite associations derived from computational microbial community models, all sources for species-metabolite correlation lay within the composition of the microbiome, while for *in vivo* species-metabolite associations causal and confounding paths linked to physiological and behavioural variation are also shaping the associating pattern.

#### Integrating in silico modelling with metabolome-microbiome association statistics

Based on these arguments, we can conclude that in the case of an *in silico* species-metabolite association, the microbiome, at least in the computational model, is causally related to the metabolite under consideration, while no conclusion about causality can be drawn for an *in vivo* association. However, as *in silico* species-metabolite associations are model-based predictions, *in silico* associations alone without empirical evidence remain hypothetical. Hence, combining metabolome microbiome association studies with COBRA community modelling holds promise for overcoming the limitations of each paradigm alone. In essence, for a given metabolite, if the species-metabolite associations *in silico* and *in vivo* systematically correlate, we can conclude that the microbiome is causally related to the metabolite and that the microbial community model is indicative of the net secretion or consumption of the metabolite through the microbiome. If no systematic correlation between *in silico* and *in vivo* association statistics is found, no conclusion can be drawn, as there are many reasons for missing correlation, from a lack of statistical power over measurement error to physiological and behavioural confounding or incomplete metabolic reconstructions of the microbes. For the same reasons, we should not expect perfect correlation between the two classes of associations, as they do not share all causal and confounding paths.

#### *In silico in vivo* association pattern analyses

These considerations lead to a methodological paradigm, called ‘*in silico in vivo* association pattern analyses’, which integrates population statistics with constraint-based modelling. This paradigm requires metagenomic data quantifying the microbiome at a body site, corresponding metabolomic measurements from the host, a collection of microbial genome-scale metabolic reconstructions, such as provided by AGORA [4, 26], for generating the microbial community models, and adequate metadata for controlling for important covariates, e.g., age, sex, and body mass index (BMI). Conceptually, *in silico in vivo* association pattern analyses can be defined by three steps (Fig 2).

1. **Step 1 (*in vivo* association pattern):** Calculate association statistics for the associations of metabolite concentrations with and the species abundances conditional on a set of covariates.
2. **Step 2 (*in silico* association pattern):** Calculate association statistics for the associations of community net metabolite secretion fluxes as determined by COBRA microbial community modelling and the species abundances conditional on a set of covariates
3. **Step 3 (*in silico in vivo* pattern analyses):** Calculate regressions for each metabolite with the significant *in vivo* metabolite-species association statistics as response variable and the corresponding *in silico* association statistics as predictor. Test the resulting regression coefficients on zero. A significant regression coefficient, and thus a significant *in silico in vivo* association pattern, indicates that the microbiome is causally related to the metabolite under consideration.

**Figure 2:**
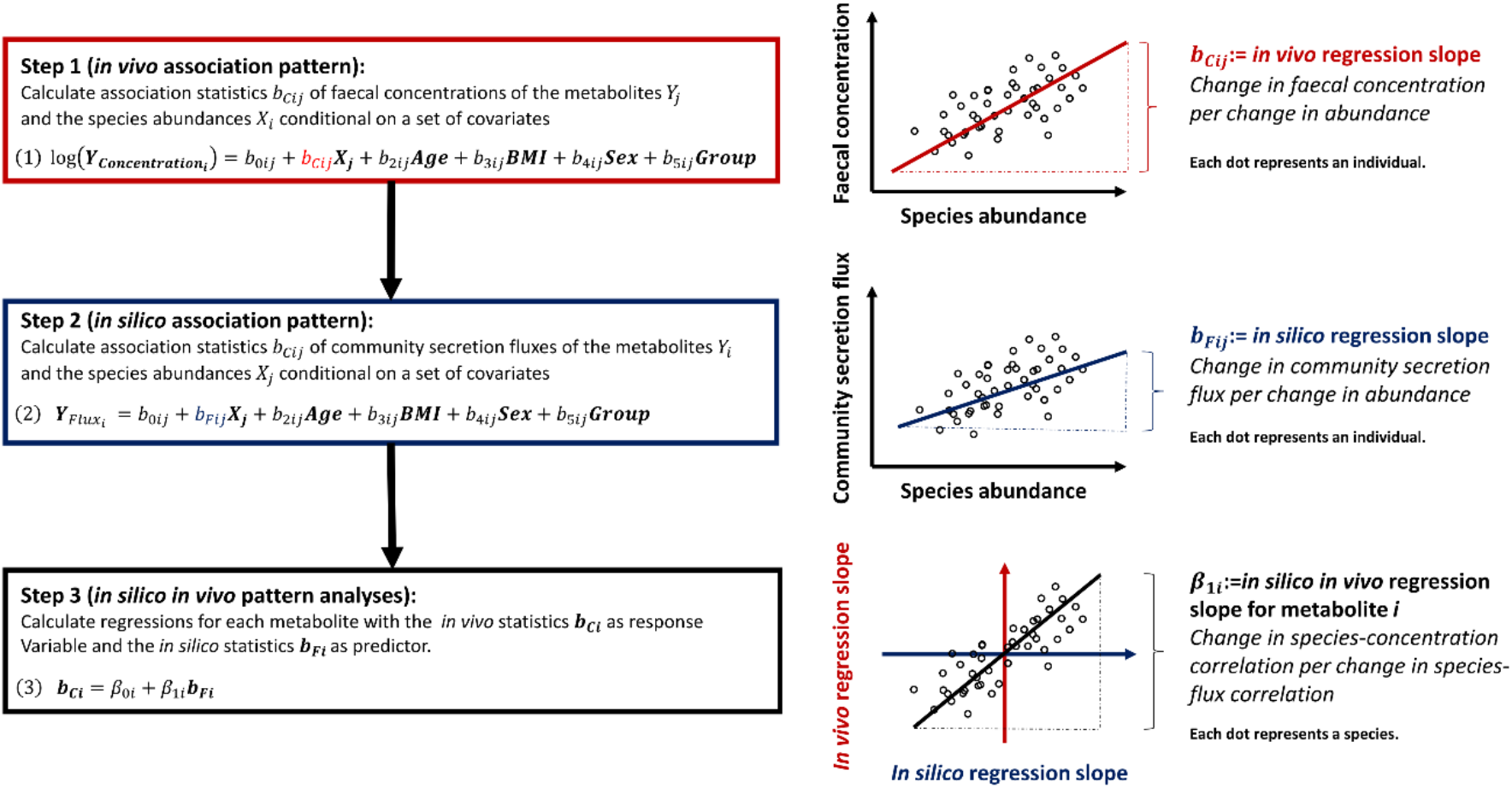
The three steps of *in silico in vivo* association pattern analyses operationalised in terms of linear regression modelling.

**Figure 3:**
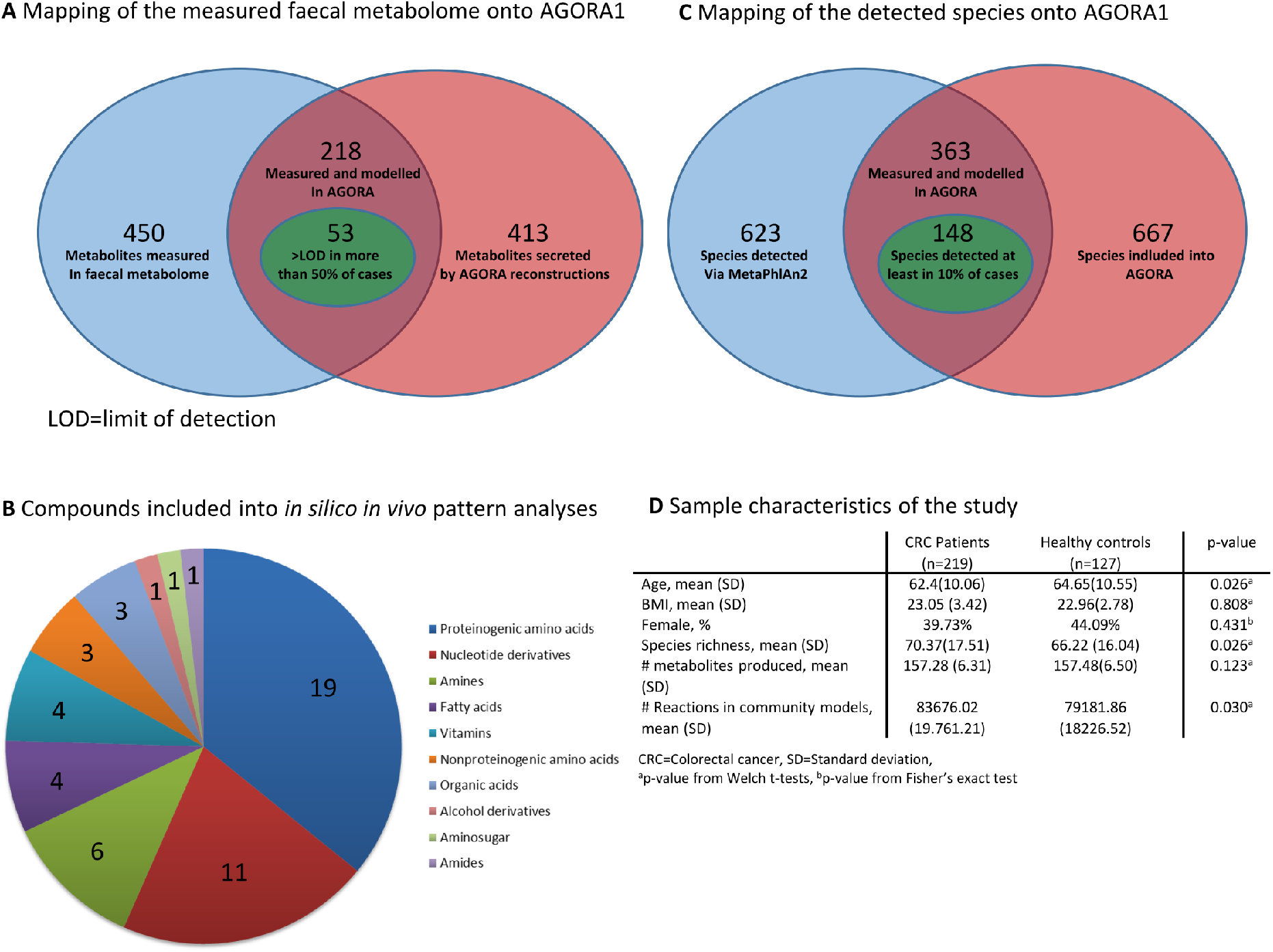
Overview on the utilised empirical dataset. **A** Mapping of the measured faecal metabolome onto the AGORA resource of gut microbial metabolic reconstructions [4]. **B** Compounds included in the *in silico in vivo* pattern analyses. **C** Mapping of the detected microbial species onto AGORA. **D** Sample characteristics of the utilised study dataset.

The concrete statistical operationalisation (i.e., which covariates to include, questions of statistical model parametrisations, and so on) is dependent on the concrete study design. The most canonical way to retrieve association statistics is via a set on linear regression models (Fig 2). However, other statistical paradigms could be used as well. For example, beyond correlating *in vivo* and *in silico* association statistics in Step 3, one may compare the sign of the two types of association statistics via simple hypergeometrical tests.

#### Summary of the theoretical part

In summary, we have established that *in vivo* species-concentration association statistics and *in silico* species-flux association statistics theoretically share certain sources of variance, while they do not completely overlap. Our theoretical considerations led consequentially to the hypothesis that for metabolites, whose concentrations in the host are systemically influenced by variance of the gut microbiome composition, we should see a substantial correlation between *in vivo* species concentration association statistics and *in silico* species flux association statistics.

This hypothesis of correlating association statistics is testable via integrating metabolome data with microbial community models based on metagenomic quantifications of the microbiome, as outlined in the section on *in silico in vivo* association pattern analysis. The most direct test can be performed by integrating computational community models of the gut microbiome with faecal metabolome data, being physiologically closest to the gut microbiome, but the outlined principles hold for other compartments and body sites as well (e.g., the oral microbiome and the saliva metabolome). Importantly, the hypothesis requires COBRA microbial community models to be fundamentally valid in their capability to predict actual metabolic activity. Testing the correlation of *in vivo* and *in silico* species-metabolite association statistics delivers a fundamental model test of COBRA community modelling regarding its ability to reflect the real metabolic activity of the gut microbiome.

### Empirical results

For testing the theoretical framework outlined above, we utilised faecal metabolome and metagenomic data from Yachida et al. [8]. First, we mapped the measured faecal metabolome data, reporting the absolute concentrations for 450 metabolites, onto the AGORA collection of 818 microbial genome-scale reconstructions [4]. We found that 106 metabolites were measured by the metabolome data and had an exchange reaction in at least one microbial reconstruction. Of these 106 metabolites, 53 had values above the limit of detection in the faecal metabolome for at least 50% of all observations and thus, they were included into the subsequent analyses (Fig 2A, 2B). Second, we mapped the relative metagenomic quantifications that had been reported by Yachida et al. [27]onto the AGORA collection. From 623 measured species, 363 were included in AGORA (2C). Next, applying personalised COBRA microbial community modelling [16], we derived for each of the 53 metabolites the net production capacity under an *in silico* average Japanese diet for 347 individuals, who had metabolome measurements (n=220 colorectal cancer cases; n=127 healthy controls). One observation from the cancer group was dropped for having zero net secretion for all metabolites due to an infeasible model configuration. Consequently, the analysis sample consisted of n=346 individuals (Fig 2D). We then calculated the *in vivo* species-concentration associations using the faecal metabolite measurements via multivariable regressions, deriving the full faecal species-concentration association pattern (Step 1). Using the *in silico* metabolic profile, we correlated the abundance of each species with the overall net community production capacity analogously (Step 2), giving rise to an *in silico* species-flux association pattern (Step 3). The species-association patterns were generated for all 148 microbial species (Fig 2C), which were found at least in 10% of all samples, resulting overall in 7844 species-metabolite associations. We generated two types of *in silico in vivo* association pattern, i) one pattern with respect to the species presence, and ii) one pattern with respect to the species abundance. For both patterns, we analysed the agreement in the sign and the value of the *in silico* and *in vivo* association statistics, resulting overall in four sets of *in silico in vivo* comparisons.

#### In vivo species presence metabolite association patterns are predicted by COBRA community models

In total, we found 2099 associations between faecal metabolite concentrations and species presence with p<0.05, and 1208 associations with a false discovery rate <0.05 (FDR) with glutarate and glutamate having the highest amount of significant species presence associations (Supplementary Table S1). For adenosine, spermidine, riboflavin, and dodecanoic acid, we found less than 10 species presence associations with p<0.05 (Supplementary Table S1). These four metabolites were dropped for missing a clear statistical *in vivo* pattern. From the remaining 49 metabolites included into the analysis, *in silico* species presence association pattern significantly predicted the sign of *in vivo* species-metabolite associations for 25 metabolites with an FDR<0.05 (Fig 4, Supplementary Table S3). Regarding the value of the *in vivo* association statistics, COBRA-based *in silico* association statistics correlated significantly (FDR<0.05) for 27 metabolites with their corresponding *in vivo* association statistics (Fig 4, Supplementary Table S3). Noteworthy, sign prediction has low statistical power if there is little variance in the signs of the *in vivo* associations (e.g., all or nearly all associations are positive, respectively negative). Still, for 23 metabolites, *in silico* associations predicted significantly (both FDR<0.05) sign and size of *in vivo* species presence metabolite associations (Fig 4).

**Figure 4:**
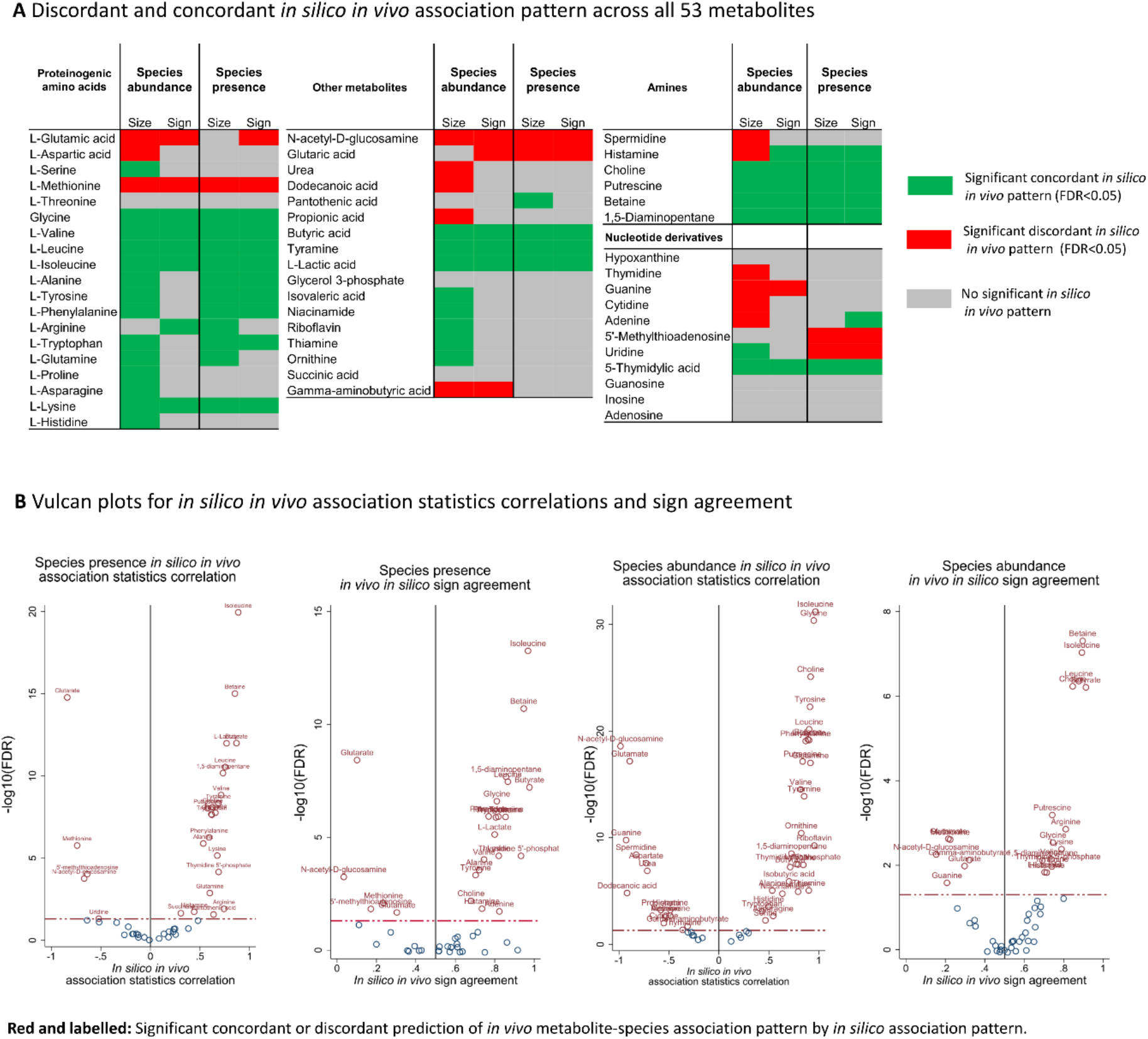
Overview on the *in vivo in silico* pattern association analyses for 53 faecal metabolites. **A** Discordance (significant, inverse association between *in vivo* and *in silico* association statistics) and concordance (significant, positive association between *in vivo* and *in silico* association statistics) for the 53 metabolites separated by metabolite class. Inconsistent pattern (uridine and thymidine) point towards model misspecifications, either in the statistical models or in the COBRA modelling. **B** Vulcan plots for the modelled metabolites displaying the strength of association (correlation and sign agreement, respectively) against the −log10 FDR. All significant patterns are labelled with the corresponding metabolites.

Importantly, for six metabolites (glutamate, methionine, N-acetyl-glucosamine, glutarate, uridine, and 5-methylthioadenosine) *in silico* associations were systematically inversed to the *in vivo* associations (Fig. 4, Supplementary Table S4). In all these cases, microbial exchange directions were noted to be bidirectional in the AGORA resource. The easiest explanation for this pattern is therefore to interpret the secretion potential as a metric of net consumption. In an earlier work [25], the validity of this interpretation was demonstrated for glutarate.

#### In vivo species abundance metabolite association patterns are predicted by COBRA community models

Calculating the *in vivo* abundance faecal concentration association pattern via regressions, we found 1305 abundance concentration associations with an FDR<0.05 and 2267 associations with p<0.05 with putrescine, alanine, and choline showing the largest numbers of species associations (Supplementary Table S2). No metabolite showed less than ten associations with at least p<0.05, and thus all metabolites were retained in the analysis. In respect to the sign of the association, the *in silico* association pattern predicted significantly the sign for 12 metabolites with an FDR<0.05 (Fig 4, Supplementary Table S4). However, in respect to the value of the association statistics, 44 out 53 metabolites showed a significant correlation between *in vivo* and *in silico* association statistics with an FDR<0.05 (Fig 4, Supplementary Table S4). As with the species presence association pattern analyses, we observed systematically inversed associations for a range of metabolites, indicating that maximal net secretion fluxes represent net consumption in these cases.

#### In silico in vivo association pattern analyses reveals broad causal microbiome-metabolome relations

Overall, *in silico* and *in vivo* association statistics significantly related to each other for 46 out of the 53 metabolites in at least one domain (i.e., value or sign for species abundance or species presence association patterns, respectively). For threonine, glycerol-3-phosphate, succinate, hypoxanthine, inosine, guanosine, and adenosine, we could not identify any significant *in silico in vivo* association pattern. Importantly, from 46 association patterns, 14 were consistently discordant (negative correlations between *in vivo* and *in silico* association statistics), 29 consistently concordant (positive correlations between *in vivo* and *in silico* association statistics), while histamine, adenine, and uridine showed mixed concordant and discordant patterns (Fig 4A, Supplementary Table S3, S4). For illustration, Figure 5 displays examples of significant and insignificant, concordant and discordant association patterns, showing the various degrees of correlations across the metabolites between *in silico* and *in vivo* association statistics. In general, all significant pattern per metabolite should be consistent, as the computed fluxes cannot indicate net consumption (discordant pattern) and net secretion (concordant pattern) at the same time. Thus, these inconsistencies hint at model misspecifications at the statistical level or at the level of COBRA community modelling, and the corresponding pattern for uridine, histamine, and adenine cannot be rated as evidence for causal relations. In conclusion, as we found clear pattern of correlation between *in silico* and *in vivo* association statistics, *in silico in vivo* association pattern analyses revealed substantial causal contributions through the gut microbiome for the faecal concentrations of 43 metabolites from a wide range of classes (Fig 4). However, *in silico in vivo* association patterns were more pronounced for amino acids and amines than for nucleotides, hinting at a higher variance contribution of the microbiome to the faecal metabolome in the domain of amino acid metabolism than in nucleotide metabolism.

**Figure 5:**
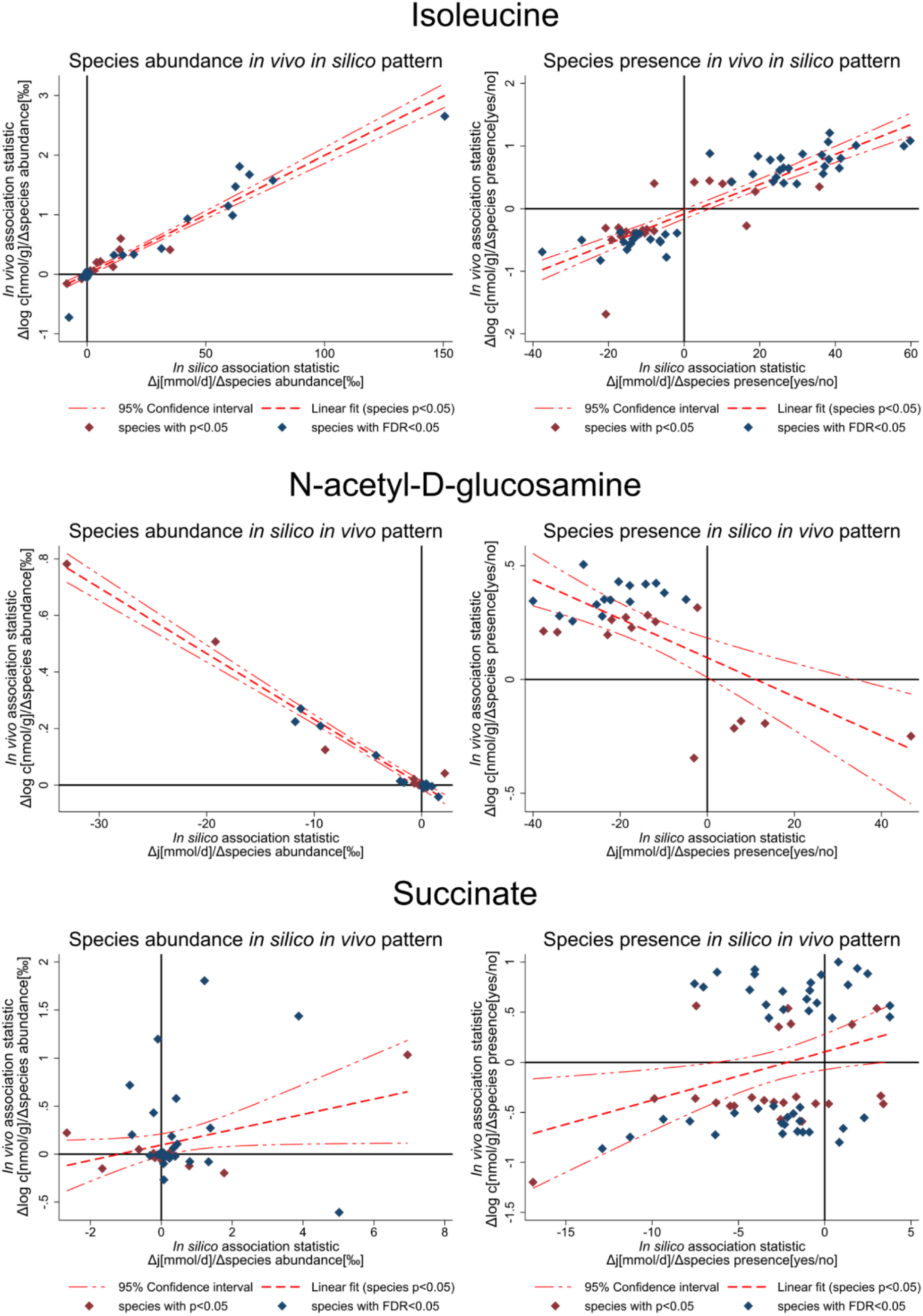
Examples for *in silico in vivo* association pattern with regression line and 95%-confidence intervals. Each dot represents a species with the X-axis denoting the *in silico* association statistic and the Y-axis denoting the *in vivo* association statistics. Association pattern for isoleucine (concordant), N-acetyl-D-glucosamine (discordant) are significant, while for succinate no significant pattern could be identified. Significance of pattern is determined by a significant slope of the regression line and significant sign (dis)agreement.

#### Prediction of the microbial component of the faecal metabolome

For certain metabolites, such as isoleucine (Fig 5), we found a very high correlation (r=0.95) between *in silico* and *in vivo* association statistics, indicating that there is a linear function between changes in community secretion fluxes and changes in log faecal concentrations. Note that this high correlation was achieved without training the community models to predict the *in vivo* associations. The empirical finding of such clear functional relationships between fluxes and concentrations indicates that it may be possible to predict the microbial component of the faecal metabolome from the microbial abundances via community modelling.

To further explore this possibility, we calculated *in silico* net concentration contributions based on the retrieved linear relationships between fluxes and concentrations from the *in silico in vivo* association pattern analyses (see Methods section for details). Therefore, we assumed that a change in species abundance, respectively species presence, would result in a change of log metabolite concentration in the faeces proportional to the corresponding change in the community net metabolite secretion flux. The latter represents a strong assumption, although the assumption seems to be plausible for those metabolites with a significant linear *in vivo in silico* association pattern.

We derived two set of prediction scores, one derived from the abundance association patterns and the second from the presence association patterns. These two prediction scores can be seen as estimates of the net concentration contribution of the microbiome to the faecal metabolome and therefore can be conceptualised as the microbial part of the faecal metabolome. The species abundance metabolite concentration prediction scores predicted significantly (FDR<0.05) the measured metabolite concentrations for 27 metabolites (Supplementary Table S5). Notably, the metabolite concentration prediction from species presence pattern was more successful with 50 metabolites being significantly predicted (FDR<0.05) and with putrescine, isoleucine, and leucine being the top hits (Fig. 6A). R-squared values reached maximally 25% in the metabolite concentration prediction from species presence pattern (Fig. 6A) and 22% in the abundance species metabolite concentration prediction, indicating that most of the variegation in the faecal metabolome is not causally related to the microbiome (Supplementary Table S5). However, the multivariate structure of the *in silico* faecal metabolite concentrations was partly reflective of the empirical correlation patterns (Fig. 6B). Especially, *in silico* faecal concentration prediction scores from species presence patterns were able to reconstruct the correlation structure for proteinogenic amino acids (Fig. 6C). Once again, the species presence metabolite prediction scores outperformed visually the species abundance metabolite prediction scores. In conclusion, *in silico in vivo* association pattern analysis allows for a characterisation of the microbial component in the faecal metabolome, partly mirroring the multivariate structure of the faecal metabolome. In this specific analysis, metabolite prediction based on species presence gave the best results.

**Figure 6:**
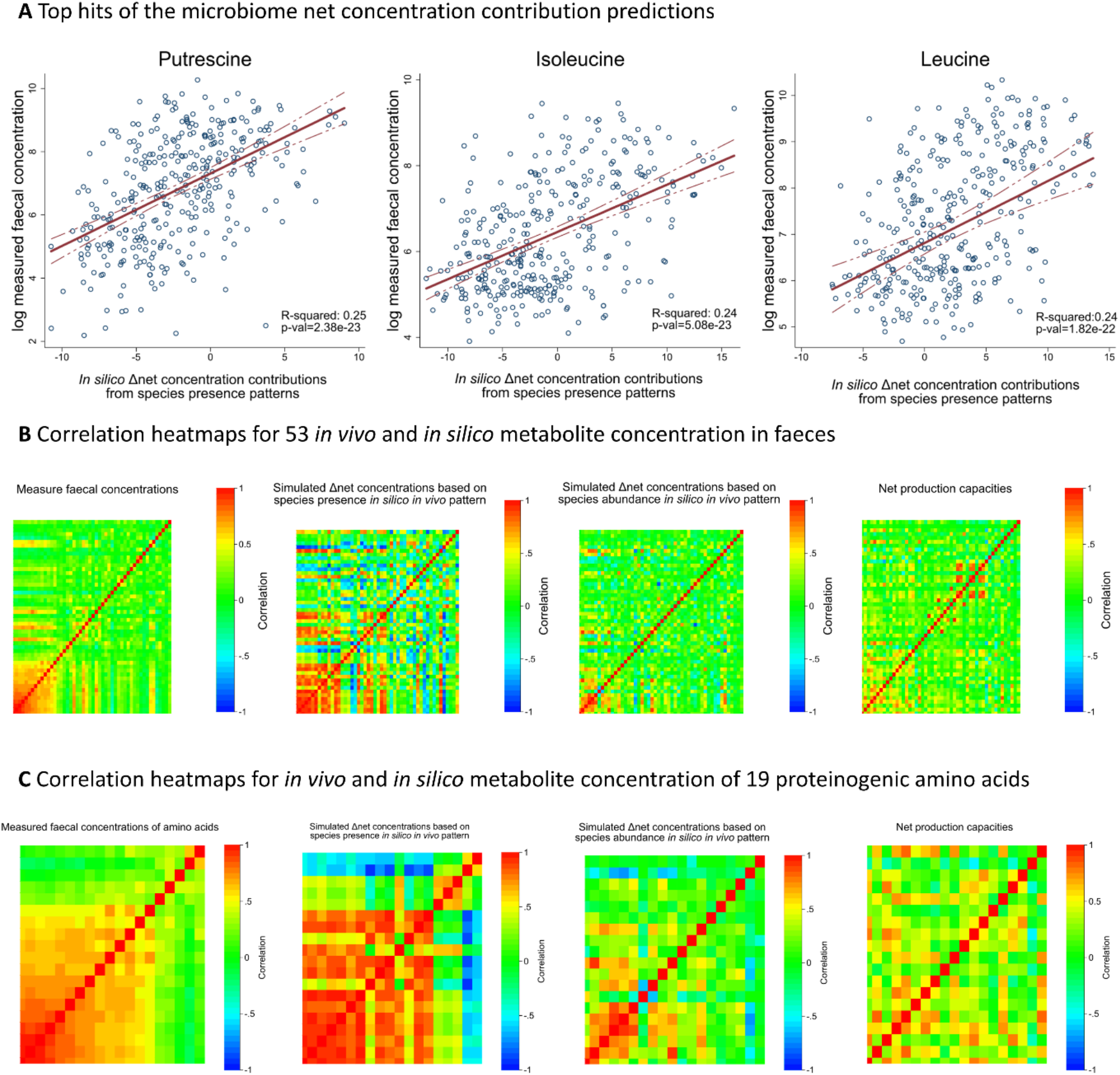
Results of the prediction of the microbial part of the faecal metabolome. **A** Scatter plots with regression lines and 95% confidence intervals for the three metabolites with highest R-squared values after prediction from the species presence *in silico in vivo* association pattern. **B** Correlation heatmaps for the 53 measured faecal metabolites, the corresponding metabolite prediction scores from species presence *in silico in vivo* association patterns, the metabolite prediction scores from species abundance *in silico in vivo* association patterns, and the raw net metabolite secretion capacities from community modelling. Rows and columns of the correlation heatmaps refer to the 53 metabolites included into modelling and the colour codes refer to the strength of the correlation between pairs of metabolites. **C** Correlation heatmaps for the 19 measured proteinogenic amino acids included into modelling, the corresponding metabolite prediction scores from species presence *in silico in vivo* association patterns, the metabolite prediction scores from species abundance *in silico in vivo* association patterns, and the raw net metabolite secretion capacities from community modelling. Rows and columns of the correlation heatmaps refer to the 19 proteinogenic amino acids included into modelling and the colour codes refer to the strength of the correlation between pairs of amino acids.

## Discussion

The microbiome contributes essentially to the human metabolism providing essential nutrients such as vitamins and short chain fatty acids, which would be inaccessible otherwise [28]. Numerous studies have been conducted to shine a light on the complex metabolic host-microbiome interplay, generating insights from a diverse range of paradigms spanning experimental work [29–31], computational and theoretical studies [11, 32] and statistical interrogation into the dependence patterns of observational multi-omics data-sets [13, 33]. Herein, we bridge the gap between population statistics and *in silico* COBRA community modelling to facilitate causal inference regarding metabolome-microbiome relations in observational data.

For identifying causal relationships in observational data, either full information on relevant covariates [34] or plausible instrumental variables (e.g., as utilised in Mendelian Randomisation [35]) must be available. In the case of microbiome-metabolome association studies, information on relevant covariates, such as diet, is typically missing and often difficult to assess reliably [13]. Moreover, while certain genetic variants have been shown to be associated with the microbiome [36], the host genetic signature is modest at best [37], making the identification of suitable genetic instrument variables to perform Mendelian Randomisation difficult. Therefore, metabolome-microbiome associations are difficult to interpret in causal terms, and indeed, simulation studies of microbial communities suggest that naïve interpretation would lead to a high level of false positives [11]. However, we demonstrate that we can overcome these difficulties by systematically integrating *in silico* COBRA microbial community modelling into statistical analyses of metabolome-microbiome datasets. On a conceptual level, COBRA microbial community modelling delivers the additional biological context needed to disentangle whether the microbiome is causally related to a metabolite or not.

Systematic *in silico in vivo* association pattern analyses revealed that most of the examined metabolites are causally related to the microbiome (81.1 %, Fig. 4). The causal signature was especially clear for amines and proteinogenic amino acids, while, relatively spoken, less pronounced for nucleotides. In a previous study, we had modelled the microbiomes of paediatric inflammatory bowel disease patients and healthy controls and predicted an increased amino acid potential in IBD microbiomes with dysbiosis that was directly linked to the presence of Gammaproteobacteria [38]. Interestingly, metabolomic measurements from the same samples had demonstrated increased faecal amino acid concentrations in IBD patients that correlated with Proteobacteria abundances [39]. While in our previous study [38], we could only indirectly compare *in silico* fluxes with metabolomic measurements, we here demonstrate the value of such an integrated analysis and showcase that the availability of multiple types of omics data for the same individuals can result in novel insight through re-analysis. The weak signature for nucleotides could indicate a higher amount of contribution of the host to the faecal nucleotide concentrations but may also point towards incomplete representation of microbial nucleotides metabolism in the genome-scale reconstructions. Importantly, the predictive power of COBRA microbial community modelling regarding *in vivo* species metabolite associations delivers a strong proof of concept that community models based on genome-scale reconstructions indeed result in quantifications of the actual metabolic activity of microbial communities.

We understand “causally related” in terms of the frameworks of Pearl [12]. It should not go unnoticed that the theory of causal statistics, built on the backbone of directed acyclic graphs, is based on strong assumptions, which can be justifiably challenged [40], especially in dynamic systems [41]. Moreover, the chosen operationalisation of causal relation can be contra-intuitive. For example, it can be that a microbial community produces a metabolite and that alterations in the community composition changes the production rate without affecting substantially the concentration in the host for various reasons (i.e., saturated transport kinetics). In this case, the microbiome would not be causal for variation in the host, while being causal for the production of the metabolite. This observation is also a reason, why *in silico* modelling alone is not sufficient for determining causal relations between metabolite concentrations in the host and species abundances. To give another example, in the case of the faecal metabolome, we only saw a weak, yet significant *in silico in vivo* association pattern regarding propionate, a short chain fatty acid known to be produced by the gut microbiome. In the case of strong intra- and interpersonal variation in absorption of propionate, which is mainly metabolised by the liver, the influence of the gut microbiome on faecal concentrations may be minor. This result indicates additionally that faecal propionate concentrations may not be good proxies for microbial propionate secretion. In contrast, *in silico in vivo* association pattern analysis was remarkably successful regarding butyrate (Supplemental Tables S3, S4), another short chain fatty acid produced by the microbiome, highlighting that faecal butyrate pools are good indicators of the microbial community butyrate production. The examples of propionate and butyrate show the value of *in vivo in silico* association pattern analyses to determine which faecal concentrations can serve as good biomarkers of microbial metabolic activity. In future studies, the influence of the host on faecal metabolite levels could be explored in simulations by integrating the microbiome models with a whole-body model of human [42].

It is worth noting that *in silico in vivo* pattern analyses, strictly spoken, allows only for an inference on whether the microbial community as a whole is causally related to a metabolite. As COBRA microbial community modelling, at least in the herein applied form, cannot differentiate between ecological causation and ecological confounding [25], causal inference on the species level, strictly spoken, is not possible. One also may argue that causal inference on single species is not sensible due to systems nature of microbial communities, making the concepts of causal inference on the species level more difficult to apply. Nevertheless, *in silico in vivo* association pattern analyses can give insights into individual species-metabolite relations. First, one can compute the direct contribution of a species to the total community’s net secretion potential [17, 38]. If the species under consideration also shows a significant species-concentration association, one may argue with some justification that a causal relationship was identified. However, one must be careful, as a species can have a positive direct contribution to the net secretion potentials, while displaying large negative ecological effects. An example of this was given in Hertel et al. [25], where *Fusobacterium* sp. were demonstrated to contribute small amounts of butyrate to faecal butyrate pools, while having large deleterious effects on community butyrate production. In this case, low abundance of prominent butyrate producers, such as *F. prausnitzii,* in *Fusobacterium* sp. containing communities resulted in an overall negative impact of *Fusobacterium* sp. on community butyrate production [25].

Other potential insights in future studies could be won by analysing outliers in the *in silico in vivo* association pattern. Outlier species (e.g., species having for example very strong *in vivo* associations, while having low *in silico* associations values) may indicate physiological and behavioural influences on metabolite-species associations not reflected in the *in silico* modelling. Thus, they could be target for further investigations in the direction of the host’s physiology and behaviour. On the other hand, outliers may also indicate incomplete genome-scale reconstructions. Thus, outlier analyses within *in silico in vivo* association pattern analysis holds promise for increasing the knowledge base and pointing towards species indicative of underlying physiological or behavioural processes.

For ten metabolites (Fig 4A), we could not identify significant or consistent *in silico in vivo* association pattern. As already sketched in the theoretical results part, the reasons for missing *in silico in vivo* association pattern can be manifold, making it impossible to disentangle the various possibilities and to conclude that the microbiome has no systematic influence on those metabolites. Table 1 categorises the different factors leading to missing association pattern into i) statistical, ii) biological, and iii) model-based factors. Importantly, we are dealing with two types of modelling (statistical modelling and COBRA modelling), each entailing their own set of assumptions. In particular, the need of sufficient statistical power to detect *in vivo* metabolite-species associations means that *in silico in vivo* association pattern analyses is bound to studies with medium to large sample sizes. Concrete sample size requirements are difficult to give, as they depend on data quality and study design, but the presented sample including 346 individuals with metabolome and metagenome data was sufficient for comprehensive analyses of *in silico in vivo* association pattern.

**Table 1:**
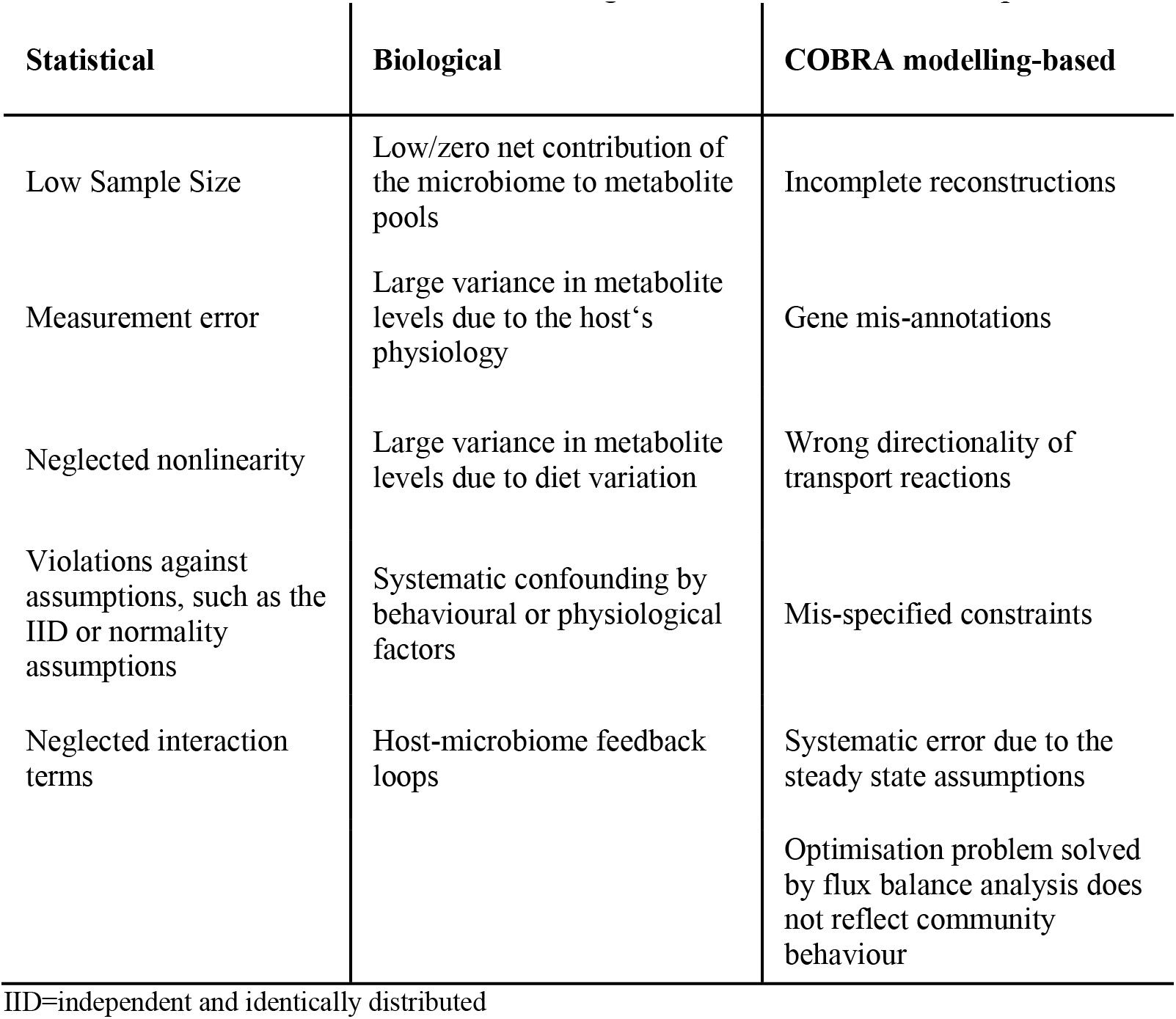
Classification of reasons for missing *in silico in vivo* association pattern

Finally, we utilised the *in silico in vivo* association pattern to derive prediction scores for faecal metabolite concentrations. The derived scores were able to reflect individual metabolite concentrations as well as mirrored the multivariate structure of the faecal metabolome. Hence, one can conclude that the microbiome is a large determining factor for the correlation structure of the faecal microbiome.

In contrast to pure machine learning approaches to predict metabolite concentrations from microbial community composition, our approach includes through genome-scale metabolic reconstructions a wealth of knowledge into the construction of the prediction scores, avoiding “conceptual overfitting” [43]. Basically, conceptual overfitting means that machine learning is blind to whether a source of covariation is due to confounding or due to causality. Maximising statistical model fit can therefore lead to parametrisations and statistical models, which may technically show the best fit, but do not approximate what we are interested in on a conceptual level in terms of biology. In the case of metabolome-microbiome studies, we are often interested in the part of the metabolome, which is causally influenced by the microbiome. As our work shows, *in silico in vivo* association pattern analyses is capable to characterise the microbial part of the faecal metabolome. In contrast, machine learning algorithms will deliver on the question, which part of the metabolome covariates with the microbiome regardless of whether the covariation is caused by confounding or causation. Notably, the *concrete* metabolome prediction scores are likely not to generalise to other human cohorts, as they are built on the covariance structure of a specific case-control study researching colorectal cancer. This problem of generalisability is common to machine learning studies and *in silico in vivo* association pattern analysis. However, applied to a large representative, general population sample, *in silico in vivo* association pattern analysis may present generalisable metabolite prediction scores, enabling to estimate the metabolic contribution of the microbiome directly from the microbial composition in a replicable manner.

Interestingly, the metabolome prediction via species presence association patterns was more successful than the prediction via species abundance association pattern. This finding is paradoxical in the first place, as species presence pattern carry less information than species abundance pattern since species are treated as binary variables (present vs. not present). The worse performance of the species abundance association pattern in predicting metabolite concentrations can have multiple reasons, both of biological and of statistical nature. While the herein applied algorithm assumes linear functions between abundances, change in fluxes, and change in log concentrations, the biologically true functions may look very different, and the functional relationships may also be species dependent. In this case, it can be that including the abundance information leads to worse performance, as it introduces substantial bias because of false parametrisations. Another methodological problem could be seen in the compositional nature of the abundance data, which is ignored in the applied prediction algorithm. Integrating compositional approaches into the prediction from *in silico in vivo* association pattern analyses may improve the performance in the future.

Regardless of future improvements, the successful prediction of metabolite concentration from association patterns shows promise for applications, where biomarkers of microbial functions are needed. For example, in pre- and probiotic interventions aiming at improving the butyrate production may be supported by such butyrate prediction scores, delivering a direct proxy of community butyrate secretion. Importantly, those scores can be superior to using butyrate quantification in the faeces, as butyrate measurements in the host are also influenced by microbiome unrelated factors, lowering arguably the statistical power to detect intervention effects.

## Conclusions

We presented a theoretical framework integrating population statistics with COBRA community modelling for causal inference on microbiome-metabolome relations. We then showed the feasibility and validity of our approach on an empirical dataset, consisting of 346 individuals with faecal metabolome data and faecal metagenomics. Conceptually, we validated thereby a methodological framework to incorporate formalised knowledge about microbial biology in the form of genome-scale models into statistical association analyses, bridging two major paradigms of systems biology. The successful identification of significant *in silico in vivo* association pattern for 43 metabolites highlights the validity of COBRA community models as theoretical model for actual microbial metabolic activity. Importantly, the prediction of *in vivo* species metabolite associations was achieved without training the COBRA community models on the given metabolome dataset. Limitations of the introduced methodology lay within the limited possibilities to perform causal inference on the level of individual species, and in the non-representation of behavioural and physiological causality, where the microbiome influences physiological or behavioural attributes of the host and thereby the metabolome. The latter aspect may be partially rectified in future studies by introducing personalised diet constraints and comprehensive whole-body modelling [42] for a more holistic picture of host-microbiome metabolic interactions. Overall, this study highlights the value of integrating knowledge-based and data-driven procedures to overcome the limitations of each paradigm alone.

## Data and code availability

All data and all scripts are freely available at https://github.com/ThieleLab/CodeBase.

## Author contributions

J.H. and I.T. designed the study. J.H. and I.T. wrote the manuscript. J.H. performed the statistical analyses and developed the theoretical concepts. A.H. constructed the microbial community models and performed the microbial community modelling. All authors reviewed and approved the final manuscript.

## Acknowledgements

This study was funded by grants from the European Research Council (ERC) under the European Union’s Horizon 2020 research and innovation programme (grant agreement No 757922) to IT, by the National Institute on Aging grants (1RF1AG058942-01 and 1U19AG063744-01).

## Methods

### Study sample

The study sample consisted of the Japanese colorectal cancer cohort data obtained from [8], which included for 347 individuals (220 colorectal cancer cases and 127 healthy controls) shot-gun sequencing data for faecal metagenomics and mass spectrometric metabolome data including quantifications for 450 metabolites. Sequencing reads and taxonomic assignments had been performed using the MetaPhlAn2 pipeline [27] Furthermore, meta-data on age, sex, and BMI were available and included into the re-analyses of the data. For details on metagenomic and metabolomic measurements, refer to [8].

### Construction of sample-specific gut microbiota models

Relative abundances on the species level for the 347 samples were obtained from the supplementary material (https://static-content.springer.com/esm/art%3A10.1038%2Fs41591-019-0458-7/MediaObjects/41591_2019_458_MOESM3_ESM.xlsx) [8]. In the first step, the quantified species were mapped onto the reference set of 818 microbial metabolic reconstructions (AGORA) [4] utilising the translateMetagenomeToAGORA.m function of the Microbiome Modelling Toolbox [16]. Next, personalised microbial community models were generated via the mgPipe module of the Microbiome Modelling Toolbox. First, pan-species models were built from AGORA version 1.03. For each metagenomic sample, the AGORA pan-species models corresponding to the species present in the metagenome were joined into one constraint-based microbial community reconstruction [16]. Next, the flux through the reactions of each pan-species model was coupled with to the flux through the respective biomass objective function (for details see [44]). Then, the community biomass reaction was parametrised via the relative abundances as stoichiometric values for each individual microbe biomass reaction. Finally, the community models were contextualised with constraints corresponding to an average Japanese Diet constraints as described previously (see below, Table S6) [16]. Finally, the community biomass reaction flux was set to be between 0.4 and 1 mmol/person/day, representing faecal excretion of once every three days to daily.

### Definition of an average Japanese diet

We used an average Japanese, as described in [26]. Briefly, the *in silico* Japanese diet was formulated based on the average daily food consumption of 106 Japanese individuals extracted from food frequency questionnaires and 28 days weighed diet records [45] (Table S6a). To convert the dietary information into constraints (given in mmol/person/day) suitable for COBRA microbial community modelling, we used the Diet Designer of the VMH database (https://vmh.life), which lists the composition of >8,000 food items [46]. In the absence of a perfect match, the closest food item entry was retrieved (Table S6a). The obtained uptake flux values were then applied to the uptake reactions present in each microbial community model using the Microbiome Modelling Toolbox [16] (see below). Ensuring the growth of all pan-species models under the defined diet, we refined the uptake fluxes as necessary (Table S6b).

### Simulations

The net community secretion flux values were determined as described in [17], [18]. Briefly, for each metabolite that could be transported by at least one AGORA model included in the community models, flux variability analysis [47] was performed for the respective dietary and faecal secretion exchanges. The net secretion fluxes, which correspond to the absolute value of the difference between the maximal flux through the faecal secretion exchange reaction and the minimal flux through the corresponding dietary uptake exchange reaction, were subsequently retrieved for each personalised model. All simulations were performed in MATLAB (Mathworks, Inc.) version R2018b with IBM CPLEX (IBM) as the linear and quadratic programming solver. The simulations were carried out using the COBRA Toolbox [15] and the Microbiome Modelling Toolbox [16].

### Statistical operationalisation of *in vivo in silico* association pattern analyses

*In silico in vivo* association pattern analyses consists of three steps. First, the *in vivo* association pattern was determined by calculating the associations between species and metabolite concentration. Second, the *in silico* association pattern between species and community net secretion fluxes was determined. Third, the pattern of *in silico* association was analysed together with the pattern of *in vivo* associations. The *in silico in vivo* association pattern analyses was performed on all 346 cases with valid COBRA community models. All statistical analysis was performed within STATA 16/MP (Stata Inc., College Station, Texas).

#### Species presence in vivo association studies

To generate the *in vivo* association pattern for species presence, we performed linear regressions with the log faecal concentration as response variable, the species presence (binary: present vs. not present) as predictor of interest, while including age, BMI, sex, and study group (binary: colorectal cancer vs. healthy controls) as covariates. To account for potential heteroscedasticity, heteroscedastic robust standard errors were used. This regression model was performed for each microbial species found in at least 10% of the samples, resulting in 148 included species, and for all metabolites having non-zero measurements in at least 50% of the cases, resulting into 53 metabolites included in the analyses. Note that, by using log transformations for the faecal concentrations, we treated zero concentration measures as missing values. We then retrieved the regression coefficient of the species presence variable, which referred in this case to the difference in mean log concentration between the individuals having a certain species in their gut microbiome and those not having this species conditional on the included vector of covariates. To assess significance, FDR correction [48] was applied correcting for 148*53=7844 tests. The regression model is given in equation (1) with 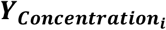 denoting the faecal concentration of metabolite *i*, 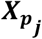 the species presence of species *j*. The regression coefficient 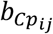 is the coefficient of interest.

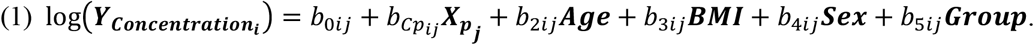

Full results can be found in the Supplementary Table S1.

#### Species presence *in silico* association studies

To derive the *in silico* species presence association pattern, community net secretion fluxes for the 53 metabolites measured in more than 50% of the cases were calculated as described above. Then, an analogous series of regressions to the *in vivo* species presence metabolite association models were performed exchanging the log faecal concentration with the net community secretion flux values. The net community secretion flux values were not log transformed, because their distributions were not consistently right-skewed as it was the case for the faecal concentrations. Then, the regression coefficients of the microbial species presence variable were extracted. The corresponding regression model is shown in (2) with 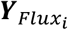 denoting the faecal concentration of metabolite *i*, 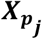 the species presence of species *j*. The regression coefficient is the coefficient of interest.

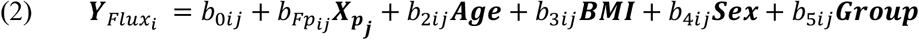

Full results can be found in the supplementary material (Table S3).

#### Species abundance in vivo association studies

To calculate the species abundance *in vivo* association pattern, we formulated analogous models to (1) by exchanging the microbial species presence variable with the species abundance. All other aspects remained the same. Full results can be found in Table S2.

#### Species abundance in silico association studies

The species presence *in silico* association pattern were derived via equation (2), exchanging the microbial species presence variable for the species abundance. Once, again all other aspects of regression modelling remained the same. Again, full results can be found in Table S4.

#### *In silico in vivo* association pattern analyses

We retrieved two pairs of association statistics (species presence pattern and species abundance pattern). To analyse the *in silico in vivo* association pattern, we calculated for each of the 53 metabolites a linear regression with the *in vivo* regression species metabolite regression coefficients as response variable and the corresponding *in silico* regression coefficient as predictor using once again heteroscedastic robust standard errors. Only microbial species-metabolite association pairs were included where the *in vivo* association statistics were at least nominally significant (p<0.05). Metabolites with less than ten nominally significant *in vivo* associations were excluded as they were missing a robust *in vivo* association pattern. The utilised regression equation is displayed in (3) and the corresponding slope β_1*i*_ was then tested on being zero. Note that ***b*_*Fi*_** and ***b*_*Ci*_** are now vectors of regression coefficients, originating from the *in silico* and *in vivo* association studies of the metabolite *i*.

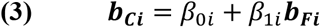

Significance of regression model (3) was determined after correction for multiple testing by using the FDR accounting for 53 tests in the case of the abundance association pattern and for 50 tests in the case of the species presence association pattern analysis.

In a second step, we analysed the agreement in sign of *in silico* and *in vivo* association statistics. This analysis was done via hypergeometrical tests, where the sign of the *in vivo* association statistics was tabulated against the sign of the *in silico* association statistics per metabolite. Once again, significance was assessed after correction for multiple testing using the FDR. Note that the statistical power to detect significant sign agreement depends on variation in the signs of the associations. Statistical power will be low if nearly all associations have the same sign.

#### Prediction of the microbial component of the faecal metabolome

We derived prediction scores for the 53 metabolites utilising the *in silico in vivo* association pattern. Two set of scores were derived: i) metabolite prediction scores based on the species presence *in silico in vivo* association pattern, and ii) metabolite prediction scores based on the species abundance *in silico in vivo* association pattern. The corresponding equations are given in (4) and (5).

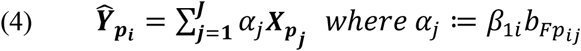

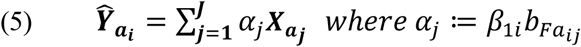

To validate the scores, we fitted linear regressions using the actual faecal concentrations as response variables and the prediction scores as predictor of interest, resulting in two sets of 53 regressions. Significance of prediction scores were evaluated after correction for multiple testing via the FDR. Additionally, we calculated the correlation matrices of the net metabolite secretion capacities, the log faecal concentrations, the metabolite prediction scores, comparing the *in vivo* correlations with the three types of *in silico* correlation matrices via correlation heatmaps.

